# JASPAR RESTful API: accessing JASPAR data from any programming language

**DOI:** 10.1101/160184

**Authors:** Aziz Khan, Anthony Mathelier

**Affiliations:** Centre for Moiecuiar Medicine Norway (NCMM), Nordic EMBL Partnership, University of Osio, 0349 Osio, Norway; Department of Cancer Genetics, Institute for Cancer Research, Osio University Hospitai Radiumhospitaiet, 0372 Osio, Norway

## Abstract

JASPAR is a widely used open-access database of curated, non-redundant transcription factor binding profiles. Currently, data from JASPAR can be retrieved as flat files or by using programming language-specific interfaces. Here, we present a programming language-independent application programming interface (API) to access JASPAR data using the Representational State Transfer (REST) architecture. The REST API enables programmatic access to JASPAR by most programming languages and returns data in seven widely used formats. Further, it provides an endpoint to infer the TF binding profile(s) likely bound by a given DNA binding domain protein sequence. Additionaly, it provides an interactive browsable interface for bioinformatics tool developers. The REST API is implemented in Python using the Django REST Framework. It is accessible at http://jaspar.genereg.net/api/ and the source code is freeiy available at https://bitbucket.org/CBGR/jaspar under GPL v3 license.

## 1 Introduction

JASPAR (http://jaspar.genereg.net) is the largest open-access database of curated transcription factor (TF) binding profiles for TFs from six different taxa (Sandelin *et al.,* 2004; Mathelier *et al.,* 2016). The underlying JASPAR data, accessible as flat files, SQL tables, or in a R data package, has been used by numerous motif enrichment tools, data analysis pipelines, and several other bioinformatics software (Stormo, 2015). Over the years, programming language-specific packages have been developed in Perl (Lenhard and Wasserman, 2002), Python (Mathelier *et al.,* 2014), R (Tan and Lenhard, 2016), and Ruby (Mathelier *et al.,* 2016) to retrieve and process data from JASPAR. However, all of these tools are language-specific and most require connection to the MySQL database.

We developed a language- and platform-independent Application Programming Interface (API) to programmatically retrieve data stored in the JASPAR database using the Representational State Transfer (REST) architecture. REST (Fielding, 2000) is a well-recognized modern web service architecture, which uses HTTP protocol to send and receive data using uniform resource locators (URLs). This new JASPAR API provides an easy-to-use RESTful web services to query/retrieve data from the latest version of the JASPAR database. The API comes with programmatic and human browsable interfaces. The HTML-based browsable interface provides a useful means for developers to interrogate the RESTful API manually before implementation in their system. From the JASPAR RESTful API, TF binding profiles can be retrieved in seven different formats: JSON, JSONP, JASPAR, MEME, PFM, TRANSFAC, and YAML.

## 2 Implementation

The JASPAR RESTful API is implemented in Python using the Django REST Framework. The source code is freely available under GPL v3 license at https://bitbucket.org/CBGR/jaspar. We used Apache HTTP server as a gateway to the API and MySQL database for persistence storage and retrieval of data requested by API services.

### 2.1 JASPAR RESTful API overview

The RESTful API is interrogated through simple URLs to provide programmatic and human browsable interface to the JASPAR database. For instance, you can access information about the TF binding matrix associated with the CTCF TF at http://jaspar.genereg.net/api/v1/matrix/MA0139.1 and get the PFM values in JASPAR format by using http://jaspar.genereg.net/api/v1/matrix/MA0139.1jaspar.

To ensure no disruption in the applications and tools using the API, the API is version-based with the version included in the API’s URL. New versions will be released with version number-based URLs, and there will be a prior notice before obsoleting any older versions. In order to control the rate of requests that clients can make to the API, we use throttling. It allows 25 requests per second from the same IP address, but no limit on the total number of requests. To enhance performance, we use Memcached (http://memcached.org), which is a fast, efficient, and fully memory-based cache server. Further, to provide a faster response to each request, the API provides pagination. The number of records per page can be modified by setting the *page_size* parameter (by default *page_size=10*). The *page* parameter is then used to request a specific page (see examples in Supplementary data). Additionally, the API supports simple query parameters to control ordering of results. For instance, one can set the parameter *order*=*name* to order TF binding profiles by name. The order can be reversed by prefixing the field name with '-' and multiple orderings may also be specified, separated by ‘,’ (see Supplementary data for examples).

The RESTful API supports several data renderer types to return responses with various media types. Currently, seven data formats are available for TF binding profiles: JSON, JSONP, JASPAR, MEME, PFM, TRANSFAC, and YAML. Users can set the output format in three different ways: (i) by setting a <*format*> parameter in the URL, (ii) by adding a <*format*> suffix to the URL, or (iii) by using the Accept headers in the get request (see Supplementary data for examples).

### 2.2 RESTful API endpoints

The RESTful API provides several endpoints to query and retrieve data from the JASPAR database (Tables 1 and S1). It also provides endpoints to infer the most likely TF binding profile(s) associated with a given protein sequence (Mathelier *et al.,* 2016). A summary of current API endpoints is available in Table 1 and a list of parameters to filter TF binding profiles from the ‘/*matrix*’ endpoint is provided in Tables 2 and S2.

**Table 1:** Summary of REST API endpoints

| Method | Endpoint | Description |
| --- | --- | --- |
| GET | /matrix | Gets a list of all profiles. * |
| GET | /matrix/:matrix_id | Detail view of the profile |
| GET | /matrix/:base_id/versions | List of profile versions |
| GET | /collection/:collection | List profiles in a collection |
| GET | /infer/:sequence | Infer profiles, given a protein sequence |

\* Profiles can be filtered using several parameters listed in Table 2 and Table S2.

**Table 2:** List of query parameters to filter profiles for API endpoint /*matrix*/

| Parameter | Description | Example |
| --- | --- | --- |
| *search* | Anything to search | ?search=Myc |
| *collection* | JASPAR collection name | ?collection=CORE |
| *tax_group* | Taxonomic group | ?tax_group=vertebrates |
| *class* | Transcription factor class | ?class=Zipper-Type |
| *family* | TF family | ?family=SMAD factors |
| *type* | Type of data/experiment | ?type=SELEX |
| *version* | Get latest matrix version | ?version=latest |

A detailed documentation for each endpoint with sample calls in multiple programming languages (JavaScript, Perl, Python, R, Java, and Ruby, following (Yates *et al.,* 2015)) is available on the API website. Further, an interactive and live API for all the API endpoints is available at http://jaspar.gene-reg.net/api/v1/live-api.

## 3 Discussion

The JASPAR RESTful API can be used to query TF binding profiles and associated data from the JASPAR database programmatically, using numerous programming languages. Moreover, the API’s web browsable interface brings an easy interface to bioinformatics tool developers.

Several third-party API clients have been developed to communicate with RESTful APIs. API clients handles the underlying details of how network requests are made and how responses are decoded. We are using Core API (http://www.coreapi.org/), which is a format-independent Document Object Model for representing web APIs. Core API comes with clients for command line, Python, and JavaScript. It is recommended to use these clients for more robust and meaningful interactions with the JASPAR RESTful API rather than constructing HTTP GET requests and decoding responses. More details about the clients are available on the API website.

The REST architecture has been demonstrated as a sustainable and reliable way to distribute genomic data to many programming languages. In the future, we plan to implement non-disruptive features to the API, such as: adding new endpoints, data fields, and filtering methods. All the disruptive changes will be released as a new version. We will make sure to keep the JASPAR RESTful API up-to-date and functional at any time to avoid any disruption to the tools based on this web service.

## Acknowledgements

We thank all JASPAR collaborators for their useful suggestions and testing the API, and Georgios Magklaras for IT support.

## API Overview

The JASPAR RESTful API provides programmatic and human-browsable access to the JASPAR database, and is implemented in Python by using the Django REST Framework.

## API Documentation

A detailed documentation of the API with sample calls in different programming languages is available at http://jas-par.genereg.net/api/vl/docs. An interactive and live API for all the API endpoints is available at http://jaspar.gene-reg.net/api/vl/live-api/.

## API Versioning

Currently, the JASPAR RESTful API has version 1, available at http://jaspar.genereg.net/api/vl. New versions will be released with new URLs with prior notice before obsoleting any older API version.

## Throttling

Our API is using throttling in order to control the rate of requests that clients can make to the API. We allow **25** requests per second from the **same IP address**, but no limit on the total number of requests.

Please feel free to contact us if you need higher request.

## Pagination

To provide a faster response to each request and to prevent larger accidental downloads, the API provides pagination. Users can increase or decrease the number of records per pages by setting the page_size parameter. By default page_size=10, which can be increased up-to **100**.

http://jaspar.genereg.net/api/v1/matrix/?page=1&page_size=25

Further, to jump from one page to another, specify the page parameter.

http://jaspar.genereg.net/api/v1/matrix/?page=2

## Ordering

The requests support a simple query parameter controlling the ordering of results. The query parameter is named order

http://jaspar.genereg.net/api/v1/matrix/?order=name

For example, to order matrices by name:

The client may also specify reverse order by prefixing the field name with ‘-’ like so:

http://jaspar.genereg.net/api/v1/matrix/?order=-name

Multiple orderings may also be specified:

http://jaspar.genereg.net/api/v1/matrix/?order=name,version

## Output formats

The JASPAR RESTful API provides several data renderer types, allowing to return responses with various media types. The query parameter is named format.

Currently, available data formats are; json, jsonp, jaspar, meme, transfac, pfm, api and yaml.

For example, to return all the matrices in **JSON** format:

http://jaspar.genereg.net/api/v1/matrix/?format=jaspar

This request will return something like this:

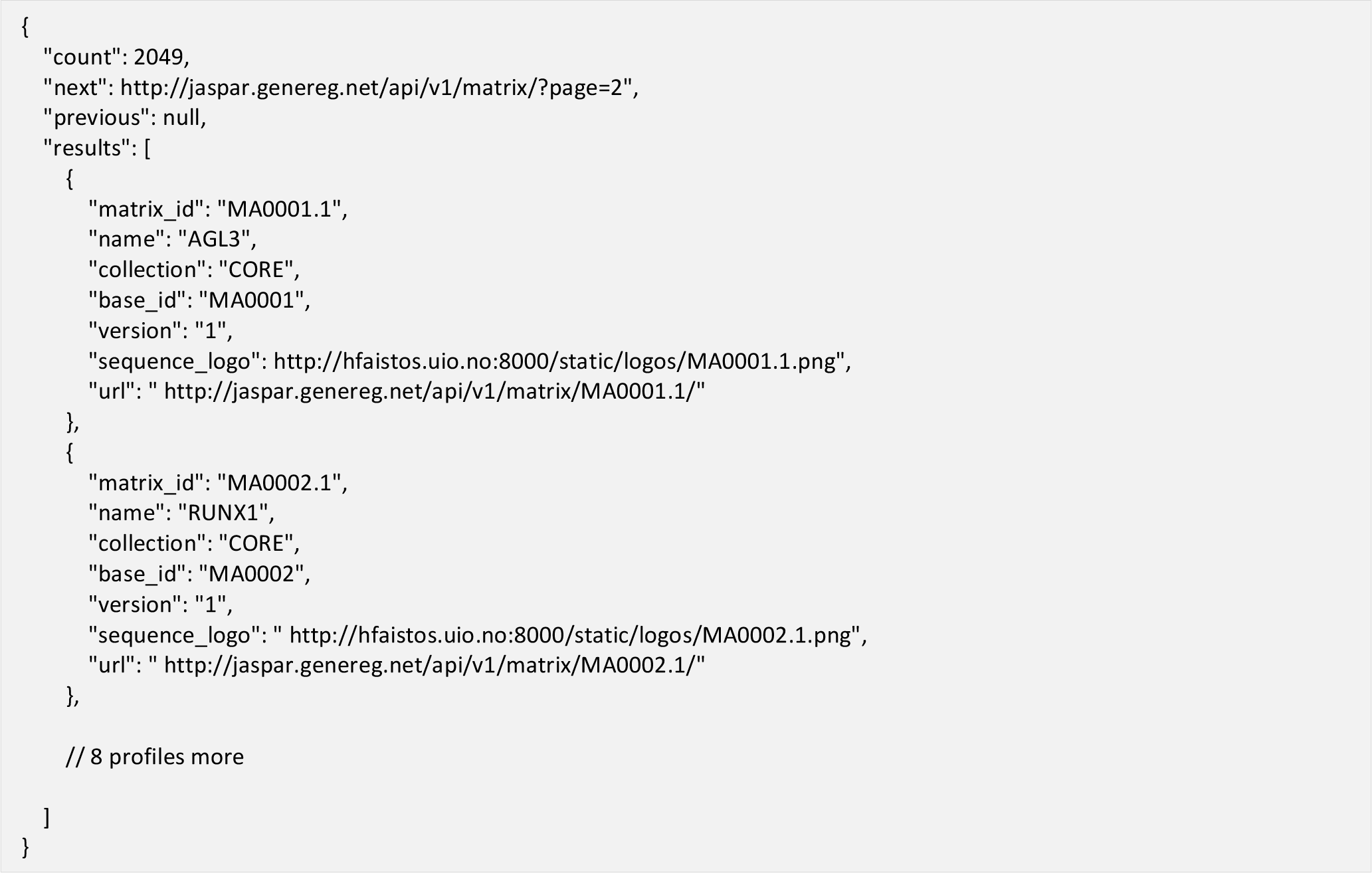

The matrix list endpoint returns multiple profiles with pagination and has the following extra fields:

- count: total number of profiles for the matching query
- next: url to get the next page profiles results. It will be null for the last page
- previous: url to get the previous page profiles results. It will be null for the first page
- results: results of the matching query.

Further, to return PFM values of matrix MA0001.1 in JASPAR format:

http://jaspar.genereg.net/api/v1/matrix/MA0001.1?format=jaspar

The above URL will return:

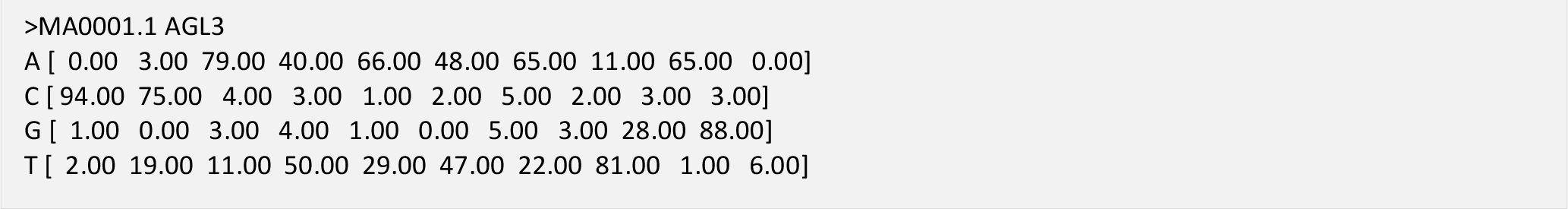

You can set the output format type in three different ways:

1. By setting ?format=format url parameter. For example:
  - http://jaspar.genereg.net/api/vl/matrix/MA0001.1/?format=json
  - http://jaspar.genereg.net/api/vl/matrix/MA0001.1/?format=jsonp
  - http://jaspar.genereg.net/api/vl/matrix/MA0001.1/?format=yaml
  - http://jaspar.genereg.net/api/vl/matrix/MA0001.1/?format=api
  - http://jaspar.genereg.net/api/vl/matrix/MA0001.1/?format=jaspar
  - http://jaspar.genereg.net/api/vl/matrix/MA0001.1/?format=meme
  - http://jaspar.genereg.net/api/vl/matrix/MA0001.1/?format=transfac
  - http://jaspar.genereg.net/api/vl/matrix/MA0001.1/?format=pfm
2. By adding .format suffix. For example:
  - http://jaspar.genereg.net/api/vl/matrix/MA0001.lJson
  - http://jaspar.genereg.net/api/vl/matrix/MA0001.lJsonp
  - http://jaspar.genereg.net/api/vl/matrix/MA0001.1.yaml
  - http://jaspar.genereg.net/api/vl/matrix/MA0001.1.api
  - http://jaspar.genereg.net/api/vl/matrix/MA0001.lJaspar
  - http://jaspar.genereg.net/api/vl/matrix/MA0001.1.meme
  - http://jaspar.genereg.net/api/vl/matrix/MA0001.1.transfac
  - http://jaspar.genereg.net/api/vl/matrix/MA0001.1.pfm
3. By using the <u>Accept headers</u> (https://www.w3.org/Protocols/rfc2616/rfc2616-secl4.html).

For example:

curl 'http://jaspar.genereg.net/api/v1/matrix/MA0001.1' -H 'Accept: application/json'

curl 'http://jaspar.genereg.net/api/v1/matrix/MA0001.1' -H 'Accept: application/javascript'

curl 'http://jaspar.genereg.net/api/v1/matrix/MA0001.1' -H 'Accept: application/yaml'

curl 'http://jaspar.genereg.net/api/v1/matrix/MA0001.1' -H 'Accept: text/html'

curl 'http://jaspar.genereg.net/api/v1/matrix/MA0001.1' -H 'Accept: text/jaspar'

curl 'http://jaspar.genereg.net/api/v1/matrix/MA0001.1' -H 'Accept: text/meme'

curl 'http://jaspar.genereg.net/api/v1/matrix/MA0001.1' -H 'Accept: text/transfac'

curl 'http://jaspar.genereg.net/api/v1/matrix/MA0001.1' -H 'Accept: text/pfm'

## API Endpoints

All the endpoints available in the current version of the API are listed in Table S1.

API ROOT: http://jaspar.genereg.net/api/v1

**Table S1:** *A detailed summary of API endpoints*

| **Method** | **Endpoint** | **Parameters** | **Description** |
| --- | --- | --- | --- |
| *GET* | /matrix | **Required:** None<br>**Optional:** All the optional parameters are listed in table S2 | Returns a list of all the matrix profiles. |
| *GET* | /matrix/:matrix_id | **Required:** matrix_id<br>**Optional:** | Gets profile detail information |
| *GET* | matrix/:base_id}/versions | **Required:** bases_id<br>**Optional:** | List matrix profile versions based on base_id |
| *GET* | collection/:collection | **Required:** collection<br>**Optional:** | Returns a list of all matrix profiles based on collection name. |
| *GET* | /infer/:sequence | **Required:** sequence<br>**Optional:** | Infer matrix profiles, given protein sequence |

**Table S2:** *Detailed list of optional parameters of API endpoint /matrix*

| Parameter | Description | Parameter Type | Data Type |
| --- | --- | --- | --- |
| page | A page number within the paginated result set. | query | integer |
| page_size | Number of results to return per page. | query | integer |
| search | Any search term. | query | string |
| order | Which field to use when ordering the results. | query | string |
| collection | JASPAR CORE or Collection name. For example: CNE | query | string |
| tax_group | Taxonomic group. For example: Vertebrates | query | string |
| class | Transcription factor class. For example: Zipper-Type | query | string |
| family | Transcription factor family. For example: SMAD factors | query | string |
| type | Type of data/experiment. For example: SELEX | query | string |
| version | If set to latest, return latest version | query | string |

## References

Fielding,R.T. (2000) Architectural Styles and the Design of Network-based Software Architectures. PhD dissertation: University of California, Irvine.

Lenhard,B. and Wasserman,W.W. (2002) TFBS: Computational framework for transcription factor binding site analysis. Bioinformatics, 18, 1135-1136.

Mathelier,A. et al. (2014) JASPAR 2014: an extensively expanded and updated open-access database of transcription factor binding profiles. Nucleic Acids Res., 42, D142-D147.

Mathelier,A. et al. (2016) JASPAR 2016: a major expansion and update of the open-access database of transcription factor binding profiles. Nucleic Acids Res., gkv1176.

Sandelin,A. et al. (2004) JASPAR: an open-access database for eukaryotic transcription factor binding profiles. Nucleic Acids Res., 32, D91-D94.

Stormo,G.D. (2015) DNA Motif Databases and Their Uses. In, Current Protocols in Bioinformatics. John Wiley & Sons, Inc.

Tan,G. and Lenhard,B. (2016) TFBSTools: an R/bioconductor package for transcription factor binding site analysis. Bioinformatics, 32, 1555-1556.

Yates,A. et al. (2015) The Ensembl REST API: Ensembl Data for Any Language. Bioinformatics, 31, 143-145.

